# Bioactivity of Aqueous Extract of Pulp of *Crysophyllum albidium* (G. Don-Holl.) on Isoproterenol Treated Albino Rats

**DOI:** 10.1101/2020.08.06.240523

**Authors:** Adetoun Morakinyo, Ifeoluwani Akinlusi, Ogundana Timileyin, Ayodeji Adepoju, Bolajoko Akinpelu, Temitope Oyedepo, Oluboade Oyedapo

## Abstract

Cardiovascular disorder is one of the causes of death globally. Quite a number of medicinal plants have been demonstrated to possess and exhibit beneficial effects on cardiovascular system throuhg their antioxidant properties. The study investigated the *in vivo* antioxidant properties of aqueous *C. albidium* pulp extract on isoproterenol-induced cardiotoxicity in albino rats. Cardiotoxicity was induced in rats by subcutaneous injection of ISO (85 mg/kg bwt) into adult albino rats and treated with the extract (100 and 200 mg/kg bwt). The levels of nitrite, lipid peroxidation, reduced glutathione total protein and activities of glutathione peroxidase, catalase and superoxide dismutase were evaluated using standard methods. The concentration of total protein was also determined. Administration of ISO (85 mg/kg body weight) caused significant (*p* < 0.05) reduction in the antioxidant status of animals. The enzymatic and non-enzymatic antioxidants perturbed by the administration of ISO were restored in the *C. albidium* treated rats. The study concludes that the aqueous extract of *C. albidum* pulp protected against Isoproterenol-induced cardiotoxicity in adult male Albino rats.

## 1. Introduction

Medicinal herbs have been used in folk medicine for millennia and in recent times, scientific study of their effects has flourished. Herbs are used by local people to treat and manage various diseases such as *Diabetes mellitus*, cancer, cardiovascular diseases amongst others [1]. The human body has a complex system of natural enzymatic and non-enzymatic antioxidant defenses which counteract the harmful effects of free radicals and other reactive oxygen species (ROS). Free radicals are responsible for causing a large number of diseases including cancer, cardiovascular diseases, neural disorders, Alzheimer’s disease mild and many metabolic disorders [2]. Protection against free radicals can be enhanced by ample intake of dietary antioxidants. Substantial evidence indicates that foods contain antioxidants and possibly, the antioxidant nutrients may be of major importance in disease prevention [3].

*C. albidum* commonly known as African star apple is one of the indigenous wild fruit tree that belongs to the family Sapotaceae and has up to 800 species whose natural occurrence is widely distributed throughout the tropical East, Central and West Africa regions particularly in Nigeria, Uganda, Niger Republic, Cameroon and Cote d’ ivoire [4]. In folklore medicine, the roots of *C. albidum* are used to treat sprains, bruises and wounds in Southern Nigeria [5, 6] while in Western Nigeria, the bark is used for the treatment of malaria and yellow fever and the leaf is used as an emollient for the treatment of skin eruption, stomachache and diarrhea [5]. African star apple (*C. albidum*) juice could protect against oxidative stress linked atherosclerosis and decrease the atherogenic index, thereby supporting the local use of *C. albidum* in the management of atherosclerosis and hypertensive conditions [7]. Hence, the present study investigated the bioactivity of aqueous *C. albidum* pulp extract on Isoproterenol induced cardiotoxicity in albino rats.

### 1.1. Materials and Method

#### 1.1.1. Plant Material: Collection and Identification

Fresh *C. albidium* fruits were obtained from a local market in Owode-Ede, Osun State, Nigeria. The fruits were identified at Adeleke University Herbarium, Department of Botany Ede, Osun State, Nigeria.

#### 1.1.2. Experimental Animals

Thirty adult male albino rats with average weight 195 ± 2.60 g were obtained from the Animal House, Ladoke Akintola University of Technology (LAUTECH) in Ogbomoso, Oyo State, Nigeria. The animals were maintained under standard laboratory conditions in the Animal House, Faculty of Science, Adeleke University, Ede, Osun State and acclimatized for 14 days. The animals were fed with standard rat pellet and watered *ad libitum*. The principle of laboratory animal care (NIH publication No 85-23) guidelines were followed in the study.

## 2. Methods

### 2.1 Preparation of Aqueous *C. albidum* Pulp Extract

The aqueuos extract of *C. albidium* was prepared as earlier reported [8]. In brief, the fleshy pulp was milled with a mechanical grinder, sieved and allowed to settle, decanted carefully and filtered. The filtrate was lyophilized to obtain the aqueous extract of *C. albidum* pulp which was stored at −20°C until further use.

#### 2.1.1 Grouping and Treatment of Animals

The experimental rats (30) were divided into five groups of six animals each and treated as follows:

Group 1 (Normal control): Rats + normal saline
Group 2 (Cardiac control): Rats + ISO (85 mg/kg)
Group 3 (Drug control): Rats + Propanolol (1.8 mg/kg bwt) + ISO (85 mg/kg)
Group 4: Rats + extract (100 mg/kg bwt) + ISO (85 mg/kg)
Group 5: Rats + extract (200 mg/kg) + ISO (85 mg/kg)

Propranolol or extract were administered orally to experimental rats, once daily for 14 consecutive days. With the exception of control rats (group I), group II-V rats were challenged with subcutaneous dose of isoproterenol at an interval of 24 hours for 2 days (14^th^ and 15^th^) to induce experimental cardiotoxicity. Animals were sacrificed 24 hours after the last dose of isoproterenol

#### 2.1.2 Collection of Blood and Heart

After the last dose of isoproterenol, the rats were fasted overnight, anesthetized with diethylether and sacrificed. The blood was collected by cardiac puncture into sample bottles containing EDTA and the heart was collected in universal bottles and stored in a refrigerator at a temperature of −4°C, for further use.

#### 2.1.3 Preparation of Heart Homogenate

The hearts were surgically removed and immediately rinsed in normal saline (0.85% w/v NaCl) to remove blood cells. The tissue (1g) was cut into thin slices and homogenized in 10 ml of freshly prepared 100 mM phosphate buffer, pH 6.8. The homogenates were centrifuged on Bench centrifuge model 800D (Pathway Medicals England, U.K) at 3000 rpm for 10 minutes. The supernatants were carefully transferred into clean vials and stored frozen for further biochemical assays.

### 2.2 Assay of the Antioxidant Potentials of *C. albidum* Extract

#### 2.2.1 Nitrite Assay

The Griess reagent was used to determine the concentration of nitrite [9]. The stock solution of sodium nitrite was prepared by dissolving 0.1 mg of nitrite in 10 ml of distilled water. The stock solution was diluted (1: 10) and further serially diluted into 5 concentrations (0 μl, 250 μl, 500 μl, 750 μl and 1000 μl) then made up to 1 ml.The samples and different standard dilutions (200 μl) and were pipetted into well labeled clean test tubes in duplicates, mixed with 1800 μl of the Griess reagent and incubated at room temperature for 10 min. The absorbance was measured at 540 nm against the blank and the standard curve was prepared by plooting absorbance of standard against concentrations. The concentration of nitrite was interpolated from the standard calibration curve and expressed in mg/ml.

#### 2.2.2. Assay of Anti-lipid Peroxidation

The lipid peroxidation assay was carried out following the modified method of [10]. The reaction mixture contained phosphate buffer (0.58 ml, 0.1 M, pH7.4), heart homogenate (0.2 ml), and ferric chloride (0.02 ml, 100 mM). The reaction mixture was incubated at 37°C in shaking water bath for 1 hour and was terminated by the addition of 1.0 ml of 10% (w/v) trichloroacetic acid (TCA) followed by the addition of 1.0 ml of 0.67% (w/v) thiobarbituric acid. The tubes were placed in boiling water bath for 20 min and then transferred to a crushed ice-bath before centrifuging at 2500 rpm for 10 min. The amount of TBARS formed was assessed by measuring the absorbance of the supernatant at 535 nm against reagent blank. The results were expressed as nM TBARS/min/mg tissue at 37°C using extinction coefficient of 1.56×10^5^ M^−1^cm^−1^.

#### 2.2.3 Assay of Glutathione Peroxidase (GPx) Activity

The assay for GPx activity was carried out according to the method of [11] based on catalytic oxidation of glutathione by hydrogen peroxide (H_2_O_2_). In the presence of H_2_O_2_, glutathione is converted to the reduced form.

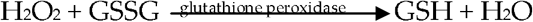

The homogenate (0.1 ml) was mixed with phosphate buffer (0.5 ml, 0.2M, pH 8.0), sodium azide (0.1 ml, 10 mM), GSH (0.2 ml, 4 mM), H_2_O_2_ (0.1 ml, 2.5 mM) and 1 ml of distilled water in duplicate. The mixture was incubated at 37°C for 3 min and TCA (0.5ml, 10%) was added. The suspension was centrifuged at 3000 rpm for 10 min followed by the collection of the supernatant. The supernatant (0.1 ml) was mixed with disodium hydrogen phosphate (0.9 ml, 0.3 M) and freshly prepared DTNB (0.6 mM, 1 ml) in sodium phosphate buffer (0.2 M, pH 8.0). The absorbance was taken at 412 nm against the blank (containing distilled water instead of the homogenate).

The GPx activity was estimated using the expression:

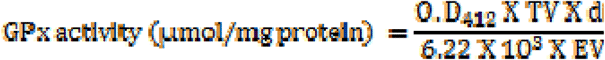

Where O.D = Absorbance at 412; df = dilution factor; extinction coefficient = 6.22 × 10^3^, EV = enzyme volume and TV = Total Volume

#### 2.2.4 Assay of Superoxide Dismutase (SOD) Activity

The assay of the heart superoxide dismutase activity was carried out according to the method of [12] based on the ability of the enzyme to inhibit the autooxidation of pyrogallol. The method’s principle is based on the competition between the pyrogallol autooxidation by O_2_•− and the dismutation of this radical by SOD.

The homogenate (200 μl) was pipetted in duplicate into clean-sterilized test tubes containing 75 mM of Tris-buffer (pH 8.2, 2.5ml), 30 mM EDTA. The reaction was initiated by the addition of pyrogallol (300 μL, 2 mM). The increase in absorbance at 420 nm was monitored every 30 secs for 150 secs against the reagent blank. The blank contained 200 μl distilled water instead of homogenate.

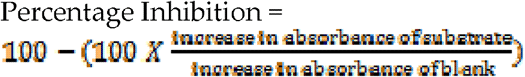

(One unit of SOD activity was given as the amount of SOD necessary to cause 50% inhibition of the oxidation of pyrogallol. The activity of SOD is expressed as Unit/mg protein.)

#### 2.2.5 Assay of Catalase Activity

The assay of heart catalase activity was carried out according to the method of [13] based on the decomposition of hydrogen peroxide to form water.

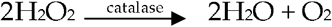

The homogenate (50 μl) in duplicate was added to a cuvette containing 450 μl of phosphate buffer (0.1 M, pH 7.4) and H_2_O_2_ (500 μl, 20 mM). Catalase activity was measured at 240 nm for 60 sec (15, 30, 45, 60 sec) against the reagent blank. The blank contained distilled water instead of the homogenate.

The catalase activity was calculated using the expression:

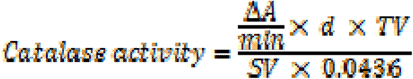

ΔA/min = slope of the graph of absorbance against min

d = dilution factor

SV = Sample volume (ml)

0.0436 = Extinction coefficient for hydrogen peroxide

TV = Total reaction volume

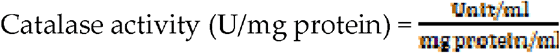

One unit of activity of catalase is equal to 1 mmol of H_2_O_2_ degraded per minute and expressed as units per milligram of protein.

#### 2.2.6 Estimation of Reduced Glutathione (GSH) Concentration

The level of reduced glutathione in the heart was estimated according to the method of [14] based on the reaction of sulfhydryl groups with DTNB (5, 5′-dithio-bis-2-nitrobenzoic acid) to produce a yellow colour, 5-thio-2-nitobenzoic acid (TNB).

Heart homogenate (1 ml) was mixed with 4 ml trichloroacetic acid 5% (w/v) and centrifuged at 4000 rpm for 10 min on a table centrifuge. The supernatant (0.1 ml) was pipetted in duplicate, 0.9 ml sodium phosphate buffer (pH 8.0, 0.2 M) and freshly prepared 5,5′-dithiobios-2-nitrobenzoic acid (DTNB) (0.6 mM, 2 ml) in 0.2 M phosphate buffer were added to it. The absorbance was read at 412 nm after 10 min against the reaction blank. Thw standard calibration curve was prepared using different concentrations of the standard GSH (0, 2, 4, 6, 8, 10 μg/ml) treated as described for the homogenate. The absorbance was plotted against the GSH concentrations. The concentration of GSH level was interpolated from the standard curve and the expressed as μg GSH/g sample.

#### 2.2.7 Estimation of Protein Concentration

Protein estimation was by the method of [15] based on the reactivity of the peptide nitrogen(s) with copper (II) ions under alkaline conditions and subsequent reduction of the Folin-Ciocateau which consists of sodium tungsate molybdate and phosphate to heteropolymolybdenum blue by copper-catalyzed oxidation of aromatic acids. Thus, the intensity of colour is proportional to the amount of these aromatic amino acids present.

The protein concentration of the test samples was extrapolated from a standard curve obtained using Bovine Serum Albumin (BSA).

## 3. Results and Discussion

Oxidative stress has been reported to play vital role in the development and progression of Myocardial infarction (MI) and heart failure [16]. There has been a direct link between activities of antioxidants and the prevention of reactive oxygen species (ROS) in ISO-administered rats [17, 18]. Studies have revealed the roles of antioxidant drugs and plant-derived compounds in the prevention of oxidative stress. In this present study, the antioxidant activity of aqueous extract of *C. albidium* was evaluated in Isoproterenol-induced cardiotoxicity with a view to studying the possibility of utilizing the plant in the management, treatment and prevention of oxidant and inflammatory related disorders. Isoproterenol is a β-adrenergic receptor agonist that causes severe stress to the myocardium of the heart muscle and produces stimulation that increases its rate and force of contraction [19].

The study showed that pre-treatment with aqueous extract of *C. albidium* was able to reduce the level of lipid peroxides significantly as compared to the cardiac control group, indicating that the extract has an anti-lipid peroxidative effect (Table 2). This may be due to the presence of phytochemicals such as flavonoids, saponins, cardiac glycosides that have been demonstrated to possess and exhibit anti-lipid peroxidation activities. Previous studies have revealed the presence of these phytochemicals [20, 8]. Numerous studies suggest that intake of fruits and vegetables that are rich in these phytochemicals have positive effects on the oxidative stress pathologies [21]. Similarly, the present study showed elevated levels of nitrite in the cardiac control group and significantly lower levels in the pre-treated groups and the reference drug group. Nitrite assay involves the reduction of nitrite to nitric oxide and other reactive nitrogen species (nitrogen oxides) using Griess reagent. These nitrogen oxides are further broken down forming stable products that are easily detected.

**Table 1:** Estimation of Nitrite and Lipid peroxidation potential.

| Group | Nitrite<br>(mg/ml) | Lipid Peroxidation<br>(nM TBARS/min/mg) |
| --- | --- | --- |
| 1 | 0.033±0.001<sup>d</sup> | 0.161±0.003<sup>c</sup> |
| 2 | 0.059±0.002<sup>a</sup> | 0.248±0.001<sup>a</sup> |
| 3 | 0.037±0.003<sup>cd</sup> | 0.144±0.005<sup>d</sup> |
| 4 | 0.044±0.001<sup>b</sup> | 0.223±0.004<sup>b</sup> |
| 5 | 0.041±0.001<sup>bc</sup> | 0.135±0.004<sup>d</sup> |

Data is represented as Mean ± SEM (n=5). Each mean value in a row was compared with the Cardiac control with superscript 'a' and different alphabet in superscript indicates a significant difference (p < 0.05) between the cardiac control and a considered value ISO: Isoproterenol

**Table 2:**
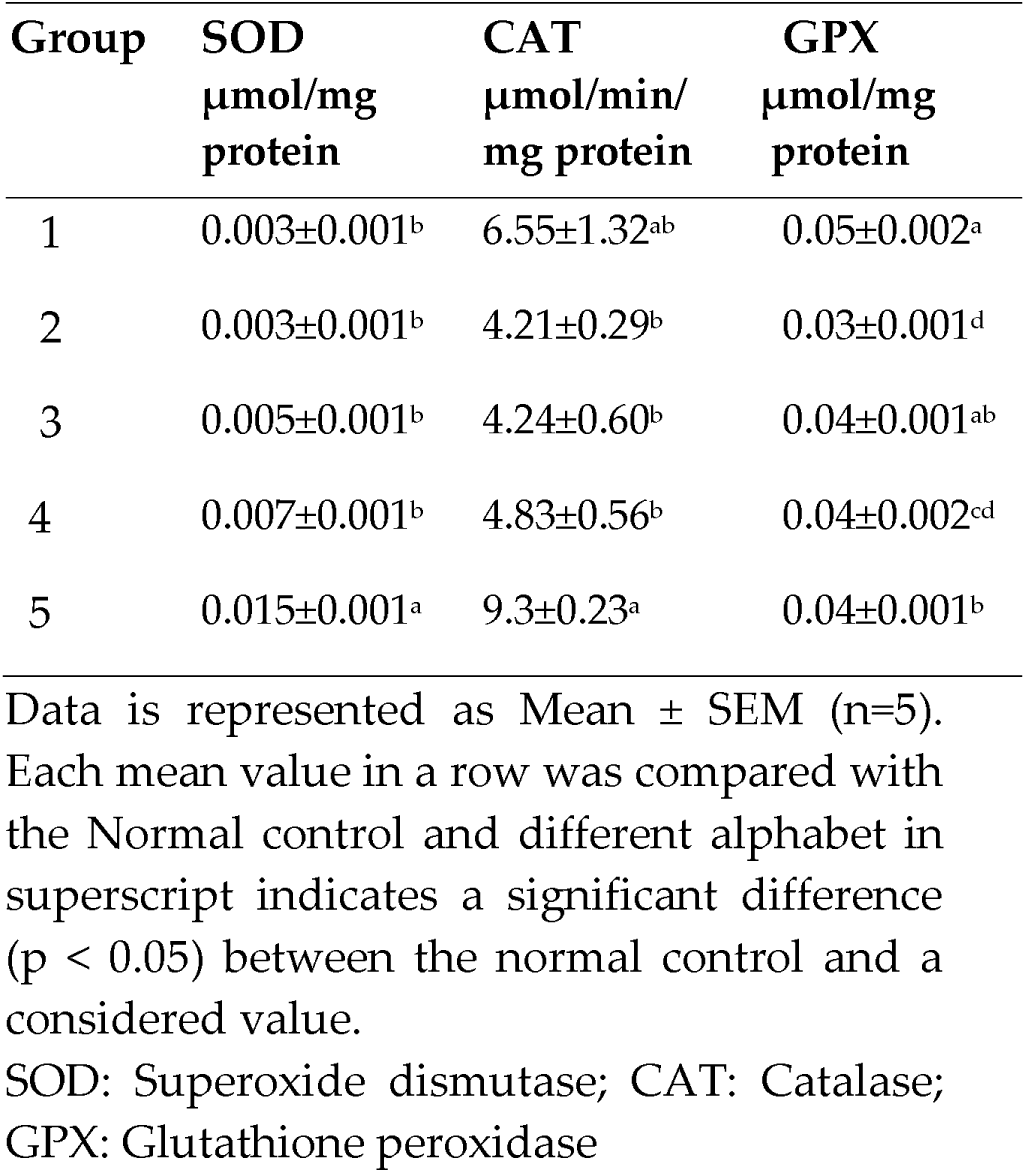
Effect of aqueous *C. albidum* pulp extract on SOD, CAT and GPx activities in Isoproterenol induced rats.

| Group | SOD<br>μmol/mg<br>protein | CAT<br>μmol/min/<br>mg protein | GPX<br>μmol/mg<br>protein |
| --- | --- | --- | --- |
| 1 | 0.003±0.001<sup>b</sup> | 6.55±1.32<sup>ab</sup> | 0.05±0.002<sup>a</sup> |
| 2 | 0.003±0.001<sup>b</sup> | 4.21±0.29<sup>b</sup> | 0.03±0.001<sup>d</sup> |
| 3 | 0.005±0.001<sup>b</sup> | 4.24±0.60<sup>b</sup> | 0.04±0.001<sup>ab</sup> |
| 4 | 0.007±0.001<sup>b</sup> | 4.83±0.56<sup>b</sup> | 0.04±0.002<sup>cd</sup> |
| 5 | 0.015±0.001<sup>a</sup> | 9.3±0.23<sup>a</sup> | 0.04±0.001<sup>b</sup> |

Data is represented as Mean ± SEM (n=5). Each mean value in a row was compared with the Normal control and different alphabet in superscript indicates a significant difference ( $p < 0.05$ ) between the normal control and a considered value.
SOD: Superoxide dismutase; CAT: Catalase; GPX: Glutathione peroxidase

The intracellular antioxidant system comprises different free radical scavenging antioxidant enzymes like superoxide dismutase (SOD), catalase and glutathione peroxidase (GPx) which constitute the first line of cellular antioxidant defense enzymes along with some non-enzyme antioxidants like reduced glutathione (GSH), vitamin C and vitamin E [22]. In Table 3 is the summary of the effect of aqueous *C. albidum* pulp extract on superoxide dismutase (SOD), catalase (CAT) and glutathione peroxidase (GPx) activities. There was no significant difference in the activities of SOD in the cardiac control rats when compared with the normal control rats. A significant increase was observed in the activities of SOD in rats treated with 200 mg/kg bwt of the extract when compared with the cardiac control rat. Superoxide dismutase (SOD), an endogenous radical scavenging antioxidant enzyme, catalyzes the removal of superoxide radicals [23] generated during ISO metabolism, to H_2_O_2_ which is a substrate for GPx and catalase enzymes. The increase of the SOD observed in this study suggests that the aqueous *C. albidum* pulp extract was involved in cellular protection and stimulation of the expression of antioxidant enzymes such as SOD [24]. The activities of catalase in the cardiac control rats reduced significantly when compared to the normal control rats. Pre-treatment with the extract at 200 mg/kg bwt caused a significant increase in the activities of catalase when compared to the cardiac control rats. However, no change was observed in the catalase activities of animals treated with propranolol and 100 mg/kg bwt *C. albidium* pulp extract treated rats when compared to the cardiac control rats. Catalase is an endogenous radical detoxifying enzyme which catalyzes the decomposition of hydrogen peroxide generated by SOD activities to water [25]. In the present study, perturbed activities of catalase were found in heart of ISO-treated rats. The aqueous *C. albidum* pulp extract probably exhibited protection against oxidative damage due to enhanced antioxidant activity and reduced the inhibition which may enhance sensitivity to free radical induced cellular damage [26]. GPx activity was significantly reduced in the cardiac control rats compared with normal control rats. Administration of the extract (100 and 200 mg/kg bwt) and propranolol increased the activities of GPx when compared to cardiac control rats. Groups 3, 4 and 5 showed similar GPX activities. Glutathione Peroxidase is a seleno-enzyme, which catalyzes the GSH dependent reduction of H_2_O_2_ and other peroxides and protects against oxidative damage [27]. According to [27], reduced glutathione peroxidase activity predicts increased cardiovascular risk following an acute coronary syndrome. In this study, the glutathione peroxidase activity of the isoproterenol only treated animals was significantly reduced when compared to the normal control animals. The increase in the GPx activity of the extract pretreated group to near normal indicated the ability of *C. albidum* pulp extract to decrease the levels of hydrogen peroxide through and maintain intracellular homeostasis as well as redox balance [18, 19, 25, 27]

There was a significant elevation in the concentration of GSH of the rats pre-treated with *C. albidium* extract (100 and 200 mg/kg bwt) when compared with cardiac control rats. Reduced glutathione (GSH) is a ubiquitous antioxidant and an essential bio-factor synthesized in all living cells, protecting the cells against free substrate for several other antioxidant radical mediated injury caused by drugs [28]. It forms important enzymes. In this study, there was a significant decrease in the reduced glutathione level of animals treated with isoproterenol only. This may be due to abundant presence of ROS in the heart of the animals and the inability of the glutathione of to restore other free radical scavengers [29]. Treatment with the extract increased the glutathione level to near normal and was dose dependent. This could be attributed to the presence of other antioxidants or free radical scavengers in the aqueous *C. albidum* pulp extract. The result of the study shows the ability of the pulp extract which possesses antioxidant properties to stabilize the integrity of cell membrane and also prevent hepatic insult mediated free radicals caused as a result of the distortion of cardiac muscles [30]. There was a significant decrease in protein concentration of cardiac control rats when compared with the normal control rats. Pre-treatment with *C. albidium* extract and propranolol increased the protein concentration when compared to the cardiac control rats. (Table 4). This suggests that an increase in protein concentration increases enzymatic activity and transport of nutrients and other biochemical compounds across cellular membranes [31].

**Table 4:** Effect of aqueous *C. albidum* pulp extract on Total Protein and Glutathione concentration in Isoproterenol induced animals.

| Group | TP<br>(mg/ml) | GSH<br>(μg GSH/g) |
| --- | --- | --- |
| 1 | 0.39 ± 0.01<sup>a</sup> | 0.04 ± 0.001<sup>a</sup> |
| 2 | 0.29 ± 0.01<sup>c</sup> | 0.03 ± 0.001<sup>c</sup> |
| 3 | 0.35 ± 0.002<sup>b</sup> | 0.04 ± 0.001<sup>b</sup> |
| 4 | 0.36 ± 0.01<sup>b</sup> | 0.04 ± 0.004<sup>b</sup> |
| 5 | 0.37 ± 0.02<sup>ab</sup> | 0.04 ± 2.800<sup>b</sup> |

Data is represented as Mean ± SEM (n=5). Each mean value in a row was compared with the Normal control with superscript 'a' and different alphabet in superscript indicates a significant difference ( $p < 0.05$ ) between the normal control and a considered value.
GSH: Glutathione; TP: Total protein

## 4. Conclusion

This study concluded that aqueous extract of *C. albidum* pulp may be considered as a cardio protective agent through the free radical scavenging mechanism. This activity may be a function of certain phytochemical constituents known to possess antioxidant properties. However, the effect is dose dependent and further studies are suggested to characterize the bioactive metabolites and its toxicity profile.

